## Supplementary materials for "Hierarchical motor adaptations negotiate failures during force field learning"

### Trajectory adaptation in Experiment-2

The PSPF is a position-dependent force field in which the force perturbes hand movements of participants predominantly in the biggining of the reach and minimally near the target. The trajectory adaptation pattern in the PSPF (Figs S1A, B and also see Fig. 1C) was similar to that in the VDCF. The PSPF perturbed the participants' hand trajectories considerably in the first adaptation and de-adaptation trials, but their hands could reach the target with the magnitude of TE being not significantly larger than the target size ($1^{st}$ adaptation trial: $t(14)=0.261$ $p=0.798$; $1^{st}$ de-adaptation trial: $t(14)=0.097$ $p=0.924$). The LD showed a monotonic change through the adaptation and de-adaptation phases although some small TEs (but larger than target size) were observed around the $2^{nd}$-$4^{th}$ adapation trials (Fig. S1A). As the TEs occurred as a consequence of over-compensation for the perturbation, their subsequent reaches might be easily adusted and little affected by the TE-driven adaptation process. Largely, the PSPF behavior followed a typical adaptation pattern with no TE and a monotonic decrease in LD, as in the VDCF, which can be explained well by the internal model adaptation alone.

The CPVF is a force field that is a combination of a position-depdendent and a velocity-dependent force fields with the two force directions opposed to each other (Fig. 1C). The velocity-dependent perturbation is predominantly effective over the first half of the reach such that the hand is pushed towards the right of the target ('+' direction). On the other hand, the position-dependent perturbation becomes stronger as the hand approaches the target, which can lead to a large TE to the left ('-' direction). This force field was utilized to comfirm that another TE-inducing force field other than the LIPF could induce the new null trajectory as observed in the Experiment-1. As we expected, large TEs were observed in the first adaptation trial (73.2 ± 50.0 (mean ± s.d.) mm, $t(13)=4.905$, $p=2.874\times10^{-4}$) as well as the first de-adaptation trial (26.0 ± 22.8 mm, $t(12)=5.211$ $p=2.178\times10^{-4}$) (Figs. S1C, D). Although some participants showed non-monotonic change of LD (see individual data in Fig S1D), the inter-participant variance of LD change was higher in CPVF compared to LIPF (Fig. 2D) such that the non-motonocity was not visible in the averaged data. This was not unexpected given that complexity of the CPVF where the velocity and position dependent part of the field push the hand in opposite directions. On the other hand, and importantly, we again observed a curved null trajectory after the de-adaptation phase of CPVF, which was clearly different from the initial null trajectory ($t(13)=3.386$, $p=0.0049$; see Fig 3). The CPVF thus could reproduce the curved null trajectory observed also after LIPF, providing further support for the presence of the TE-driven adaptaion process.

The the flat models (internal model adapation only) could again not reproduce the CPVF behaviors (Figs. S2A, B). The flat models inevitably produce a monotonic decrease in the LD in the initial trials and reduce it to zero after de-adaptation. In contrast, the hierarchical models (Fig. S2C) could again explain the CPVF behavior. The simulation again showed the necessity of the TE-driven adapation process in addition to the internal model adaptation to explain behaviors in the presence of faliures, or TEs.

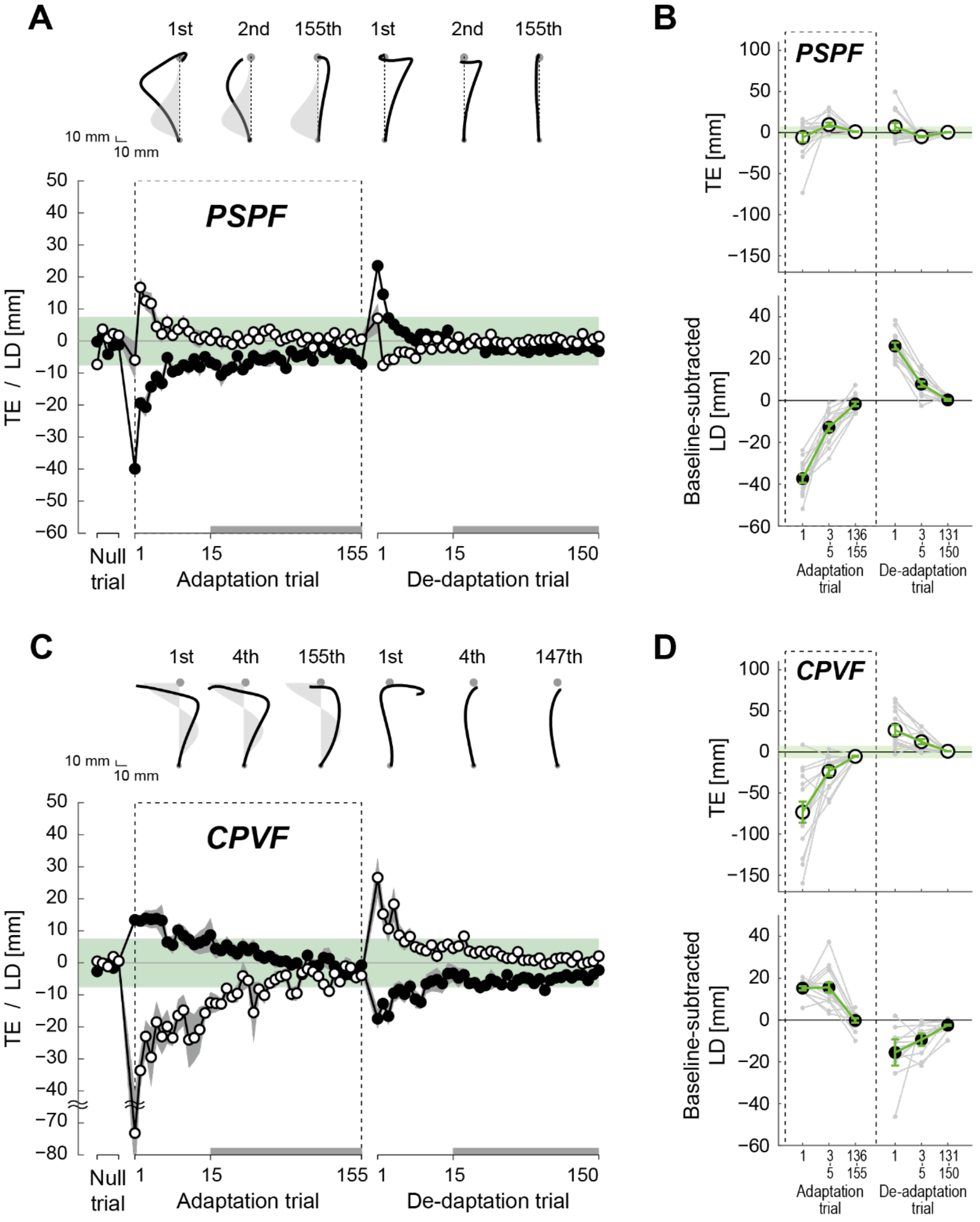


*S1. Trajectory adaptation in Experiment-2: The learning curves in PSPF (A) and CPVF (B)*

*averaged across all participants. Note that the scales differ between x and y axes to clearly show trajectory changes along the x-axis. The adaptation of the TE and LD are shown by traces with open circles and filled circles, respectively. The first 15 TE and LD values are plotted for every single trial, while the subsequent trials (indicated by thick gray lines at the bottom of the figure) are plotted for every five trials. The shaded gray areas around the lines represent standard errors. The light green zones represent the target width (radius: 7.5 mm).*

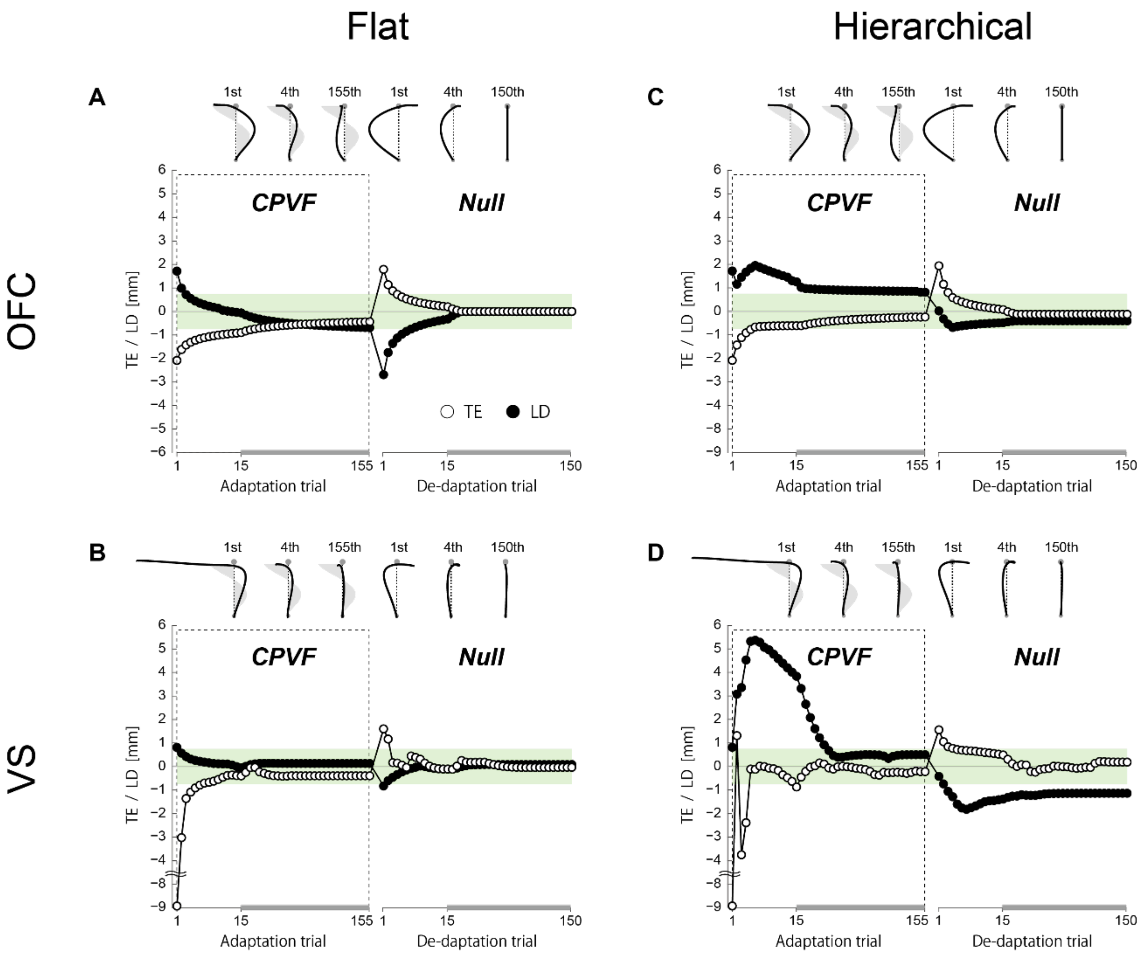


*S2. Simulation results for trajectory adaptation in CPVF, represented by TE (open circle) and LD (filled circle) by the flat (A, B)/hierarchical (C, D) OFC (upper panels) and V-shaped models (lower panels). The flat learning models (only internal model adaptation) were unable to reproduce either the non-monotonic change in LD or the curved null trajectory with a persistent deviation after exposure to LIPF. However, the hierarchical OFC models (kinematic plan learning and internal model adaptation) successfully reproduced both.*

### Detail for the simulation

To explain adaptive behaviors in the VDCF and the LIPF of the Experiment-1, we utilized two motor learning models: one is proposed by Izawa et al. (2008), which we refer to as the flat OFC model, and the other is proposed by Franklin et al. (2008), which we refer to as the flat VS model. These original models implement only the internal model learning and can explain monotonic trajectory adaptation as observed in the VDCF. However, they cannot explain non-monotonic trajectory adaptation, nor a persistent change in the null trajectory in the LIPF. We thus extended the two models by introducing a TE-driven kinematic plan adaptation that hierarchically interacts with the internal model adaptation (Fig. 6A). We referred to the extended models as the hierarchical OFC model and the hierarchical VS model, respectively.

*OFC model*

Stochastic optimal feedback control (OFC) has been extensively used to model reaching (Izawa et al., 2008, Mistry et al., 2013, Todorov and Jordan, 2002, Todorov, 2005, Guigon et al., 2008, Cesonis and Franklin, 2020). The OFC model assumes a linear dynamical system represented in a discrete-time formulation as:

$$\mathbf{x}_{t+1} = \mathbf{A}\mathbf{x}_t + \mathbf{B}\mathbf{u}_t + \xi_t + s_c \sum_{i=1}^{c} \varepsilon_t^i \mathbf{C}_i \mathbf{u}_t \qquad (1)$$

where $\mathbf{x}_t$ is the state of the system at time $t$; $\mathbf{u}_t$ is the control signal input to the system; $\mathbf{A}$ is the state transition matrix; $\mathbf{B}$ is the control input matrix; $s_c$ is the scaling factor, $\mathbf{C}_i$ are constant matrices, and $\varepsilon_t^i$ is standard normal gaussian noise for control-dependent noise; $\xi_t$ is zero-mean Gaussian noise. The model assumes a partially observable system given by:

$$\mathbf{y}_t = \mathbf{H}\mathbf{x}_t + \omega_t \qquad (2)$$

where $\mathbf{y}_t$ is the observation made by the system; $\mathbf{H}$ is the observation matrix; $\omega_t$ is zero-mean Gaussian noise. The system is assumed to have the goal of minimizing a cost over a movement. The cost accrued at each time step is given by:

$$\mathbf{x}_t^T \mathbf{Q}_t \mathbf{x}_t + \mathbf{u}_t^T \mathbf{R} \mathbf{u}_t \qquad (3)$$

where the control cost matrix $R$ is symmetric positive semidefinite ($\mathbf{R}>0$); the state cost matrix $\mathbf{Q}_t$ is symmetric positive semidefinite ($\mathbf{Q}_t \geq 0$). Under these assumptions, Todorov's method (2005) provides a solution to calculate the optimal controller and estimator as follows:

$$\mathbf{u}_t = -\mathbf{L}_t \hat{\mathbf{x}}_t \qquad (4)$$

$$\hat{\mathbf{x}}_{t+1} = (\mathbf{A} - \mathbf{B}\mathbf{L}_t)\hat{\mathbf{x}}_t + \mathbf{K}_t(\mathbf{y}_t - \mathbf{H}\hat{\mathbf{x}}_t) + \eta_t \qquad (5)$$

where $\hat{\mathbf{x}}_t$ is the state estimate of the system at time $t$; $\mathbf{L}_t$ is the optimal control gain; $\mathbf{K}_t$ is the optimal estimator gain or Kalman gain; $\eta_t$ is zero-mean Gaussian noise. $\mathbf{L}$ and $\mathbf{K}$ are calculated recursively to minimize the expected summation of cost in Eq.3. Please see Todorov (2005) for further details.

Similarly to Izawa et al. (2008), we modeled the arm reaching as a model of control for a point mass in the Cartesian coordinates. The inertia was $\mathbf{m}=\begin{bmatrix}1 & 0\\ 0 & 1\end{bmatrix}$ (kg). The state was defined as an 8-dimensional vector as follows:

$$\mathbf{x}_t=[\mathbf{p}_t^T,\mathbf{v}_t^T,\mathbf{f}_t^T,\mathbf{T}_t^T]^T \qquad (6)$$

where $\mathbf{p}_t$ and $\mathbf{v}_t$ are the 2-dimensional position and velocity of the arm, respectively; $\mathbf{f}_t$ is the 2-dimensional force produced by the arm; $\mathbf{T}_t$ is the 2-dimensional target position. As the simulation is done for the VDCF or LIPF condition, the external force imposing to the arm is written by the form:

$$D\mathbf{x}_t=\mathbf{K}_f\begin{bmatrix}0 & -1\\ 0 & 0\end{bmatrix}\mathbf{p}_t+\mathbf{B}_f\begin{bmatrix}0 & -1\\ 1 & 0\end{bmatrix}\mathbf{v}_t \qquad (7)$$

where $D\mathbf{x}_t$ is state-dependent force; $\mathbf{K}_f$ and $\mathbf{B}_f$ are coefficients for position-dependent and velocity-force fields, respectively, as such $\mathbf{K}_1$ and $\mathbf{B}_1$ (see the section of force fields in Methods). The relationship between the forces and the control signals was modeled as a first-order linear system with a time constant of $\tau$=120 ms, similar to Izawa et al. (2008). The system dynamics discretized with a time step of $\Delta$=10 ms was defined as follows:

$$\mathbf{p}_{t+1}=\mathbf{p}_t+\mathbf{v}_t\Delta t \qquad (8)$$

$$\mathbf{v}_{t+1}=\mathbf{v}_t+\left(\frac{\mathbf{f}_t+D\mathbf{x}_t}{\mathbf{m}}\right)\Delta t \qquad (9)$$

$$\mathbf{f}_{t+1}=\mathbf{f}_t\left(1-\frac{\Delta t}{\tau}\right)+\mathbf{u}_t(1+\sigma_c\varepsilon_t)\frac{\Delta t}{\tau} \qquad (10)$$

The cost was as follows:

$$\begin{array}{ll} w_r(u_x^2+u_y^2) & (0\le t<T)\\ w_p[(p_x-T_x)^2+(p_y-T_y)^2)]+w_v(v_x^2+v_y^2)+w_r(u_x^2+u_y^2) & (MT\le t<MT+MT_H)\end{array} \qquad (11)$$

where the parameters, $w_p$, $w_v$, and $w_r$, are weights for target accuracy, terminal velocity, and control signal input, respectively; $MT$ is the maximum movement completion time; $MT_H$ is the time for which the hand was supposed to hold a position at the target after movement completion.

The original model (i.e., flat OFC model) by Izawa et al. (2008) utilizes OFC to simulate reaching trajectories during adaptation to a state-dependent novel force field, based on a concept that motor learning is a process to acquire a model of the novel environment and use the model to re-optimize movements. Accordingly, in this framework, motor adaptation is characterized by the knowledge of the environment (the novel force field) which the motor system gradually acquires. The external force imposing to the arm is written by $\mathbf{D}\mathbf{x}_t$ (Eq. 7). What the motor system needs to perform the optimal movement in the force field is the full knowledge of **D**, which is assumed to be gradually acquired. The knowledge of **D** during adaptation is thus represented by the form:

$$\hat{\mathbf{D}} = \alpha \mathbf{D} \qquad (12)$$

where $\hat{\mathbf{D}}$ is the estimated force matrix, and $\alpha$ is the learning parameter, which is assumed to gradually increase from 0 to 1 with adaptation. Accordingly, during adaptation, the motor system produces the motor command optimized for the environment where the predicted external force defined as $\hat{\mathbf{D}}\hat{\mathbf{x}}_t$ could impose to the arm. Only when $\alpha = 1$, does the system have the full knowledge of **D** and produce the optimal motor commands for the actual environment. When $0 < \alpha < 1$, the system has an incomplete knowledge of **D** and would produce a sub-optimal movement for the actual environment. Thus, by changing the value of $\alpha$, Izawa et al. (2008) simulated reaching trajectories in several phases of motor adaptation.

For the hierarchical OFC model, we borrowed the idea of a kinematic bias of movement direction proposed by Mistry et al. (2013), which we refer to as directional bias. Mistry et al. extended the cost function of OFC by including a directional bias to explain a directional preference of reaching trajectories observed during motor adaptation to an acceleration-based force field. The directional bias represents the desired direction of movement, which is represented by the form:

$$\mathbf{Q}_d = \begin{bmatrix} d_y^2 & -d_x d_y \\ -d_x d_y & d_x^2 \end{bmatrix} \qquad (13)$$

where $\mathbf{Q}_d$ is the directional bias matrix and $\mathbf{d} = \begin{bmatrix} d_x & d_y \end{bmatrix}^T$ is the desired directional vector represented as a unit vector. The new terms related to the directional bias (third term in Eq. 14) were added to the original cost function (first and second terms) so that any position or velocity perpendicular to the desired direction was penalized as follows:

$$\mathbf{x}_t^T \mathbf{Q}_t \mathbf{x}_t + \mathbf{u}_t^T \mathbf{R} \mathbf{u}_t + e^{-t/\tau_d} (k_p \mathbf{p}_t^T \mathbf{Q}_d \mathbf{p}_t + k_v \mathbf{v}_t^T \mathbf{Q}_d \mathbf{v}_t) \qquad (14)$$

where $k_p$ and $k_v$ are the weight of bias for position and velocity, respectively. The exponential decay term is included because the directional bias need not exist for the entire motion. In our simulation, these parameters were set as follows: $k_p = k_v = 0.5$ , $\tau_d = 130$ ms. The cost parameters included in **Q** and **R** (Eqs. 3 and 11) were set as follows: $w_p = 10^{-7}, w_v = 0.1, w_f = 0, w_p = 30$. *MT* and *MT*$_H$ were set to 400 ms and 50 ms, respectively. We performed the simulation only considering the sensory noise, $\omega_t$ (Eq. 2) and control-dependent noise (Firth term in Eq. 1) for simplicity. Hence, the variance of zero-mean Gaussian noises of $\xi_t$ (Eq. 1), and $\eta_t$ (Eq. 5) was set to zero. The observation matrix (Eq. 2) was formulated so that the system was able to observe hand positions, velocities, and the target positions. Thus, the observation by the system was as follows:

$$\mathbf{y}_t = [\mathbf{p}_t^T, \mathbf{v}_t^T, \mathbf{T}_t^T]^T + \omega_t \qquad (15)$$

The vector $\omega_t$ is zero-mean Gaussian noise with diagonal covariance:

$$\mathbf{\Omega}^{\omega} = (diag[np, np, nv, nv, nt, nt])^2 \qquad (16)$$

where *np*, *nv*, and *nt* are the sensory noise for position, velocity, and target, respectively. *np*, *nv*, and *nt* was set to 0.05, 0.05, and 0, respectively. The multiplicative control-dependent noise added to the control signal (Eq. 1) is as follows:

$$s_c \sum_{i=1}^{c} \varepsilon_t^i \mathbf{C}_i \mathbf{u}_t = s_c \begin{bmatrix} \varepsilon_t^1 & \varepsilon_t^2 \\ -\varepsilon_t^2 & \varepsilon_t^1 \end{bmatrix} \mathbf{u}_t \qquad (17)$$

We chose c = 2 with $\mathbf{C}_1 = \begin{bmatrix} 1 & 0 \\ 0 & 1 \end{bmatrix}$ and $\mathbf{C}_2 = \begin{bmatrix} 0 & 1 \\ -1 & 0 \end{bmatrix}$ similarly to Todorov and Jordan (2002) or Guigon et al. (2008). Multiplying $\mathbf{u}_t$ by $\begin{bmatrix} \varepsilon_t^1 & \varepsilon_t^2 \\ -\varepsilon_t^2 & \varepsilon_t^1 \end{bmatrix}$ produces 2D Gaussian noise with circular covariance, whose standard deviation is equal to the length of the vector $\mathbf{u}_t$ (Todorov and Jordan, 2002). The scaling factor, $s_c$ was set to 0.01.

Here, we further extended this idea by introducing a directional bias modulated by trial-by-trial TE (upper panel, Fig. 6B). The directional bias is inclined in the opposite direction of TE to reduce it. The direction of the directional bias in the *i*-th trial is represented by $\varphi(i)$, the angle from the target direction (clockwise as positive). The TE is equivalent to the directional error represented by $\theta^i$, defined as the angle between the target direction from the start position and the direction from the start position to the endpoint of the reaching. In the presence of TE (i.e., TE > target size), the directional bias is updated according to the directional error as follows:

$$\varphi^{i+1} = b\varphi^i - r\theta^i \qquad (18)$$

where the constant *b* is the forgetting rate and is set to 0.95. The constant *r* is the sensitivity to the degree of the directional bias update to the directional error and set to 0.85. The initial value of the directional bias is 0 (i.e. $\varphi(1)=0$ ).

In the absence of TE (i.e., TE < target size), we assumed that the direction bias subtly decays across trials to the original direction towards the target as follows:

$$\varphi^{i+1}=b\varphi^{i} \qquad (19)$$

Additionally, we assume that the kinematic plan adaptation is also affected by the motor cost of the generated reaching, which is defined as $J=\int_{t=0}^{MT_H}[\mathbf{x}_t^T\mathbf{Q}_t\mathbf{x}_t+\mathbf{u}_t^T\mathbf{R}\mathbf{u}_t+e^{-t/\tau}(k_p\mathbf{p}_t^T\mathbf{Q}_\mathbf{d}\mathbf{p}_t+k_v\mathbf{v}_t^T\mathbf{Q}_t\mathbf{v}_t)]dt$. The decay of the directional bias stops, i.e., $b$ = 1 when $J$ goes below less than 0.01. The threshold value was arbitrarily determined to produce curved null trajectories similar to those in the experiments. Once a TE greater than the target size occurs, the kinematic bias is active. In contrast, if the TEs keep within the target size throughout the experiment, the kinematic bias remains inactive.

To simulate the internal model adaptation in novel force fields, we changed the value of learning rate $\alpha$. In the adaptation phase, $\alpha$ is increased from 0 to 0.8 such that $\alpha^i=0.8\cdot\log(\log(i)+1)/\log(\log(155)+1)$ for $1\le i\le 155$. In the de-adaptation phase, $\alpha$ is decreased from 0.8 to 0 in the first 30 de-adaptation trials because de-adaptation process is well known to be much faster than adaptation process (Shadmehr and Wise, 2005). This was given by $\alpha^i=0.8\cdot\log(\log(i)+1)/\log(\log(155)+1)$ for $156\le i\le 185$ ; $\alpha(i)=0$ for $186\le i\le 305$. The movement distance was 150 mm. The reach duration was set to 400 ms. For the simulation of VDCF and LIPF, $B_1$ and $K_1$ were set to 7 Ns/m and 120 N/m, respectively. For the CPVF, $B_1$ and $K_1$ were set to 9 Ns/m and 75 N/m. PEC was applied over the second half of movement (y > 7.5 cm) as a one-dimensional spring force (1500 N/m) and damper (100 Ns/m) along x-axis. These parameters were chosen to produce trajectories similar to those in the experiments.

*V-shaped model*

The original model (i.e., flat VS model) assumes that desired trajectory, which the motor system should trace, is a fixed straight line joining the start and target and that motor command is gradually corrected to reduce the difference between the actual and desired trajectory, which is defined as movement error. The VS model uses a 2-joint 6-muscle arm model to simulate the reaching trajectories in a broad range of novel force field environments.

The dynamics of the arm model is described in joint space by the form:

$$\boldsymbol{\tau}_{RB}(\mathbf{q},\dot{\mathbf{q}},\ddot{\mathbf{q}})+\mathbf{J}^T(\mathbf{q})\mathbf{F}_E=\mathbf{J}_m^T\mathbf{m} \qquad (20)$$

where $\boldsymbol{\tau}_{RB} = (\tau_s, \tau_e)^T$ are torques at the shoulder and elbow joints; $\mathbf{F}_E = (F_x, F_y)^T$ are the forces exerted on the hand in the Cartesian coordinate; $\mathbf{m}$ are muscle tensions. The forces are transformed into joint torques using the Jacobian:

$$\mathbf{J}(\mathbf{q}) = \begin{bmatrix} -l_s s_s - l_e s_{se} & -l_e s_{se} \\ l_s s_s + l_e s_{se} & l_e c_{se} \end{bmatrix} \qquad (21)$$

where $s_s \equiv \sin q_s, s_{se} \equiv \sin(q_s + q_e), c_s \equiv \cos q_s, c_{se} \equiv \cos(q_s + q_e)$ . $\boldsymbol{\tau}_{RB}(\mathbf{q}, \dot{\mathbf{q}}, \ddot{\mathbf{q}})$ are the dynamics due to inertial and velocity-dependent forces, where $\mathbf{q} = (q_s, q_e)^T$ are the vector of shoulder and elbow joints. The dynamics of the 2-joint 6-muscle arm model can be written as follows:

$$\begin{aligned} &\boldsymbol{\tau}_{RB} = \Omega(\mathbf{q}, \dot{\mathbf{q}}, \ddot{\mathbf{q}})\mathbf{p} \\ &\Omega_{11} = \Omega_{21} = \ddot{q}_s + \ddot{q}_e, \quad \Omega_{12} = c_e \ddot{q}_e - s_e \dot{q}_e (2\dot{q}_e - \dot{q}_e), \\ &\Omega_{22} = c_e(\ddot{q}_s + \ddot{q}_e) + s_e \dot{q}_s^2, \quad \Omega_{13} = 0, \quad \Omega_{23} = \ddot{q}_e, \\ &p_1 = I_e + M_e l_{m,e}^2, \quad p_2 = M_e l_s l_{m,e}, \quad p_3 = I_s + M_s l_{m,s}^2 + M_e l_s^2, \end{aligned} \qquad (22)$$

where $M_s$ and $M_e$ are the masses of the upper arm and lower arm, respectively; $l_s$ and $l_e$ are the corresponding segment lengths; $l_{m,s}$ and $l_{m,e}$ are the distances to the respective centers of the mass of the segments, and $I_s$ and $I_e$ are the respective moments of inertia. The vector of muscle tensions is defined as:

$$\mathbf{m} = (m_{s+}, m_{s-}, m_{e+}, m_{e-}, m_{b+}, m_{b-})^T \qquad (23)$$

where $m_s, m_e, \text{and } m_b$ are the muscle tension of shoulder, elbow, and biarticular muscles, respectively; the subscripts '+' and '-' indicates the flexor and extensor muscles, respectively. These muscle tensions are transformed into joint torques using the Jacobian $\mathbf{J}_m(\boldsymbol{\rho})$, which is a constant matrix consisting of the muscle moment arms $\boldsymbol{\rho} = (\rho_{e+}, \rho_{e-}, \rho_{bs+}, \rho_{bs-}, \rho_{be+}, \rho_{be-})^T$ :

$$\begin{bmatrix} \tau_s \\ \tau_e \end{bmatrix} = \begin{bmatrix} \rho_{s+} & 0 \\ -\rho_{s-} & 0 \\ 0 & \rho_{e+} \\ 0 & -\rho_{e-} \\ \rho_{bs+} & \rho_{be-} \\ -\rho_{bs+} & -\rho_{be-} \end{bmatrix}^T \begin{bmatrix} m_{s+} \\ m_{s-} \\ m_{e+} \\ m_{e-} \\ m_{b+} \\ m_{b-} \end{bmatrix} \qquad (24)$$

where $\rho_s, \rho_e, \rho_{bs}, \text{and } \rho_{be}$ are the moment arms of shoulder muscles, elbow muscles, biarticular muscles around the shoulder, and biarticular muscles around the elbow, respectively; the subscripts '+' and '-' indicates the flexor and extensor muscles, respectively.

For each muscle, the tension is assumed to depend on the motor command *u*, muscle length $\lambda$, and the change rate of length $\dot{\lambda}$ as follows:

$$m = m(\lambda, \dot{\lambda}, u) \qquad (25)$$

Muscle tension consisting of two terms is given by:

$$m = \left[m_A(u) + m_{IMP}\right]_+, \quad [\cdot]_+ \equiv \max\{\cdot, 0\} \qquad (26)$$

where $m_A(u)$ is due to muscle activation $u$; $m_{IMP}$ is due to mechanical impedance produced by the muscle (i.e. muscle stiffness and damping). This muscle impedance is given by the form:

$$m_{IMP} = \kappa(E(t) + \kappa_d \dot{E}(t)) \qquad (27)$$

where $\kappa$ is muscle stiffness; $\kappa_d$ is the ratio of muscle viscosity to stiffness; *E* is the movement error. The error, *E*, is represented in coordinates of muscle length and written by the form:

$$E = \lambda - \lambda_0 \qquad (28)$$

where *E* is the difference between the actual muscle length, $\lambda$ and the desired muscle length, $\lambda_0$. The intrinsic stiffness $\kappa$ is modeled to increase linearly with the total motor command $w$ as follows:

$$\kappa(t) = \kappa_0 + \kappa_1 w(t) \qquad (29)$$

$w$ consists of the feedforward motor command, inherent motor noise, $u_N$, and neural feedback or reflex $v$ as follows:

$$w = \left[u + u_N + v\right]_+ \qquad (30)$$

The overall effect of the many different sources of variance is modeled as noise in the motor command:

$$u_N(t) = (\mu_0 + \mu_1 u)\mu(t) \qquad (31)$$

where $\mu(t)$ is 0 mean Brownian motion. The neural feedback or reflex is given by the form:

$$\mathbf{v}_t = r(E(t-\phi) + r_d \dot{E}(t-\phi)) \qquad (32)$$

where $\phi$ is the feedback delay. The model assumes that muscle tension is equal to the motor command as follows:

$$m_A = w = \left[u + u_N + v\right]_+ \qquad (33)$$

The feedforward command *u* for each muscle *k* is updated from trial *i* to trial *i+1* according to the following learning law:

$$\begin{aligned} & u^{i+1}(t) \equiv [u^i(t) + \Delta u^i(t+\phi)]_+, \ [\cdot]_+ \equiv \max\{\cdot, 0\} \\ & \Delta u^i(t) = \alpha\varepsilon_+^i(t) + \beta\varepsilon_-^i(t) - \gamma, \ [\cdot]_- \equiv [-,\cdot]_+ \\ & \varepsilon^i(t) = E^i(t) + g_d \dot{E}_k^i(t) \end{aligned} \qquad (34)$$

where $E(t)$ is the muscle stretching/shortening at time *t* (Eq. 28), and $\Delta u$ is phase advanced by $\phi > 0$, which is equal to the feedback delay. The superscript *i* represents the trial number, $\alpha$ and $\beta$ are the learning parameters ($\alpha > \beta > 0$) and $\gamma$ (>0) is a constant de-activation parameter. The term $g_d$ (>0) indicates the relative level of velocity error to length error.

Here, we extend the flat VS model by introducing an idea that the desired trajectory (lower panel in Fig. 6b), which is represented in the Cartesian coordinates, is updated according to a trial-by-trial TE in a similar way to the hierarchical OFC model. The desired trajectory is described as a curved line with a deflection, *dx*, 120 mm away from the start position along the y-axis (Fig. 6B). Before adaptation, the desired trajectory is the straight line towards the target, that is, *dx* = 0. In the presence of TE (i.e., TE > target size), *dx* is updated as follows:

$$dx^{i+1} = b \cdot dx^{i} - r \cdot TE^{i} \qquad (35)$$

where the constant *b* represents the retention of motor learning and is set to 0.95; the constant *r* to the degree of update of *dx* to the TE in the previous trial and is set to 0.45. The constant *r* is thus the sensitivity to the degree of the desired trajectory update to TE. In the presence of endpoint error, *dx* is modulated such that the desired trajectory is deflected in the opposite direction to a trial-by-trial TE. The desired trajectory with *dx* was calculated as the minimum jerk trajectory with the via-point at [dx 120] (mm) from the start position (Flash & Hogan, 1985).

In the absence of TE (i.e., TE < target size), we assumed that the desired trajectory subtly decays across trials to the original direction towards the target as follows:

$$dx^{i+1} = b \cdot dx^{i} \qquad (36)$$

We again assume that the kinematic plan adaptation is affected by the motor cost of the generated reaching, which was calculated as average muscle tension across all the 6 muscles during movement. When the cost goes below less than 500, the decay of the desired trajectory stops, i.e., *b* = 1. The threshold value was again arbitrarily determined to produce curved null trajectories similar to those in the experiments.

In simulation, the desired trajectory was converted from the Cartesian to muscle space via inverse kinematics and it was applied to the learning law (Eq. 34). The trajectory of the desired muscle length $\lambda_0$ (Eq. 28) was calculated from the minimum jerk trajectory with the via-point at [dx 0]. The kinematic parameters were as shown in Table S1.

Table S1: Anthropometric data for arm segments

| | Mass (kg) | Length (m) | Center of mass from proximal joint (m) | Mass moment of inertia (Kg m$^2$) |
|---|---|---|---|---|
| Upper arm | 1.93 | 0.31 | 0.165 | 0.0141 |
| Forearm | 1.52 | 0.34 | 0.19 | 0.0188 |

The moment arms (cm) are as follows: shoulder monoarticulars: $\rho_{s+} = \rho_{s-} = 3.0$ ; elbow monoarticulars: $\rho_{e+} = \rho_{e-} = 2.1$ ; shoulder biarticulars: $\rho_{bs+} = \rho_{bs-} = 4.4$ ; elbow biarticulars: $\rho_{be+} = \rho_{be-} = 3.38$. The noise parameters (Eq. 31) are as follows:

$$\mu_0 = 7,\ \mu_1 = 0.04,\ \mu_0 = 12.5 f(v) \qquad (37)$$

where $v$ is a standard normal random variable; $f(\cdot)$ is a causal fifth-order Butterworth filter with 2Hz cut-off frequency. The delay parameter $\phi$ was set to 60 ms. The ratios of damping to stiffness for muscle and feedback components were set to $\kappa_d = 1/12$ s and $r_d = 2$ s, respectively. The two parameters for the intrinsic stiffness (Eq. 29) were set to $\kappa_0 = 3360$ Nm$^{-1}$ and $\kappa_1 = 118$ m$^{-1}$. The parameter $r$ for the neural feedback or reflex (Eq. 32) was set to 336 Nm$^{-1}$.

Similarly to Franklin et al. (2008) and Tee et al. (2010), the muscle tensions of the elbow muscles were modeled by the form:

$$m_{A,el\pm} = \left[u_{el\pm} + u_{N,el\pm} + v_{el\pm}\right]_+ + 0.3 m_{A,bi\pm} \qquad (38)$$

For the learning parameters, $g_d$ (Eq. 34) was set to 0.2. $\gamma$ (Eq. 34) was given by:

$$\gamma = \frac{2\varepsilon_{ss}\alpha\beta}{\alpha + \beta} \qquad (38)$$
$$\beta = 0.7\alpha$$

where $\varepsilon_{ss} = 7.8 \times 10^{-4}$ m; $\alpha = 9800$. Please see Franklin et al. (2008) and Tee et al. (2010) for the rationale for the parameter selection. For simulation of the arm model, we used the same parameters as those in the original studies (Franklin et al., 2008).

Finally, the parameters for the experimental environment were set as follows. The start and target positions were at [0, 350] and [0, 500] (mm) in the Cartesian coordinate (where [0, 0] is at the shoulder joint), respectively. The reach duration was set to 400 ms. For simplicity, all noise parameters were set to zero. For the force fields of VDCF and LIPF, $B_1$ and $K_1$ (see the section of *force field*) were set to 20 Ns/m and 120 N/m, respectively. For the CPVF, $B_1$ and $K_1$ were set to 14 Ns/m and 100 N/m, respectively. PEC was applied over the second half of movement (y > 7.5 cm) as a one-dimensional spring force (2500 N/m) and damper (1000 Ns/m) along x-axis. These parameters were chosen to produce trajectories similar to those in the experiments. Similarly to our experiments in which the hand motion was constrained to the final hand position, only for the TE-inducing force fields (e.g. LIPF and CPVF) and the Null field after the LIPF adaptation, the constrain force was applied with a strong stiff two-dimensional spring force (1000 N/m) and damper (500 Ns/m) when the y-axis hand velocity fell below 20 mm/s, which allowed us to reproduce large target errors. The simulation was completed when the hand velocity fell below 20 mm/s.
